# Sperm DNA damage causes genomic instability in early embryonic development

**DOI:** 10.1101/681296

**Authors:** Sjors Middelkamp, Helena T.A. van Tol, Diana C.J. Spierings, Sander Boymans, Victor Guryev, Bernard A.J. Roelen, Peter M. Lansdorp, Edwin Cuppen, Ewart W. Kuijk

**Affiliations:** Center for Molecular Medicine and Oncode Institute, University Medical Center Utrecht, Utrecht University, Universiteitsweg 100, 3584, CG, Utrecht, The Netherlands; Department of Farm Animal Health, Faculty of Veterinary Medicine, Utrecht University, Yalelaan 104, Utrecht 3584 CM, The Netherlands; European Research Institute for the Biology of Ageing, University of Groningen, University Medical Center Groningen, Groningen 9713 AV, the Netherlands; Department of Medical Genetics, University of British Columbia, Vancouver, BC, V6T 1Z4, Canada; Terry Fox Laboratory, BC Cancer Agency, Vancouver, BC, V5Z 1L3, Canada

## Abstract

Genomic instability is common in early embryo development, but the underlying causes are largely unknown. Here we examined the consequences of sperm DNA damage on the embryonic genome by single-cell genome sequencing of individual blastomeres from bovine embryos produced with sperm damaged by radiation. Sperm DNA damage caused fragmentation of chromosomes and segregation errors such as heterogoneic cell divisions yielding a broad spectrum of genomic aberrations that are similar to those frequently found in human embryos. The mosaic aneuploidies, mixoploidy, uniparental disomies and *de novo* structural variation induced upon sperm DNA damage may compromise health and lead to rare genomic disorders when embryos escape developmental arrest.

**One Sentence Summary:** DNA damage in sperm cells leads to genomic defects in embryos

## Main Text

In early embryonic development, there is reduced activity of cell cycle checkpoints and apoptotic pathways until the zygotic genome becomes activated (*1-4*). As a consequence, mitotic errors are tolerated in the first cleavage divisions causing genomic instability and mosaicism, i.e. the phenomenon that cleavage stage embryos are composed of multiple genetic lineages. Mosaicism affects approximately three quarters of the human day 3 cleavage stage embryos and contributes to the low success rate of in vitro fertilization (IVF) through high miscarriage rates and failed implantations (*5-7*). On rare occasions, chromosomally abnormal cells may develop to molar pregnancies or contribute to parthenogenetic, androgenetic chimaeric, and mixoploid lineages in live-born humans (*8-12*). Mosaicism is prevalent in human spontaneous abortions of natural pregnancies (*13, 14*), indicating that the causes for the high mitotic error rate in embryos are unrelated to the IVF procedures such as the ovarian stimulation regime, fluctuations in oxygen tension or temperature, and composition of the culture medium. While advanced maternal age increases the risk for meiotic errors leading to whole embryo aneuploidies, mitotic errors and embryo mosaicism are not correlated with female age (*15-17*). A minor allele of the *polo-like kinase 4* (*PLK4*) gene has been associated with tripolar chromosome segregations, leading to mitotic errors in development (*16, 17*). However, *PLK4* polymorphisms alone cannot explain the high prevalence of mosaicism in human embryos. Thus, the causes for the high mitotic error rates in human preimplantation embryos are still largely unknown (*6, 18, 19*).

The role of the sperm cell in embryonic mosaicism has thus far been largely ignored (*20*), possibly because paternal effects on the embryonic genome are presumed to be mostly restricted to the zygote stage. Here, we used bovine IVF to investigate the consequences of sperm DNA damage on embryonic genome integrity. Bovine IVF is recognized as a valuable model system to study genomic instability in mammalian embryos (*21, 22*).

Subjecting sperm cells to increasing doses of γ-radiation reduced blastocyst formation rates (Fig. S1A) and caused developmental arrest at around the eight-cell stage (*4*), which coincides with the activation of the zygotic genome (*23*). Development up to the eight-cell stage thus appears to be a deterministic process regulated by maternally deposited factors that support the first cleavage divisions irrespective of the degree of DNA damage to the sperm cell. This provides a window of opportunity to naively study the role of sperm DNA damage on genomic instability in the absence of selection.

To study the consequences of sperm DNA damage on the embryonic genome we performed Strand-seq, a single-cell genome sequencing technique in which the DNA strands that were used as templates during DNA replication prior to cell division are selectively sequenced. Strand-seq enables the genome-wide detection of copy neutral and copy number structural variants and strand-inheritance patterns between daughter cells (*24-26*). Strand-seq libraries display the characteristic Watson-Watson, Watson-Crick, or Crick-Crick patterns of the template strand that was inherited by the daughter cell after cell division (*24*). Strand-seq was performed on both blastomeres of two-cell stage embryos (∼28 hours post-fertilization (hpf)) produced with sperm subjected to zero (0 Gy), a low (2.5 Gy) or high (10 Gy) dose of radiation. The sequenced libraries of 47 individual blastomeres derived from 26 two-cell stage control embryos and 73 individual blastomeres of 47 two-cell stage embryos produced with damaged sperm that passed quality control were further analyzed (Fig. S1B,C, Table S1). Sister cells displayed the expected complementary strand inheritance patterns (Fig. 1A).

**Fig. 1.**
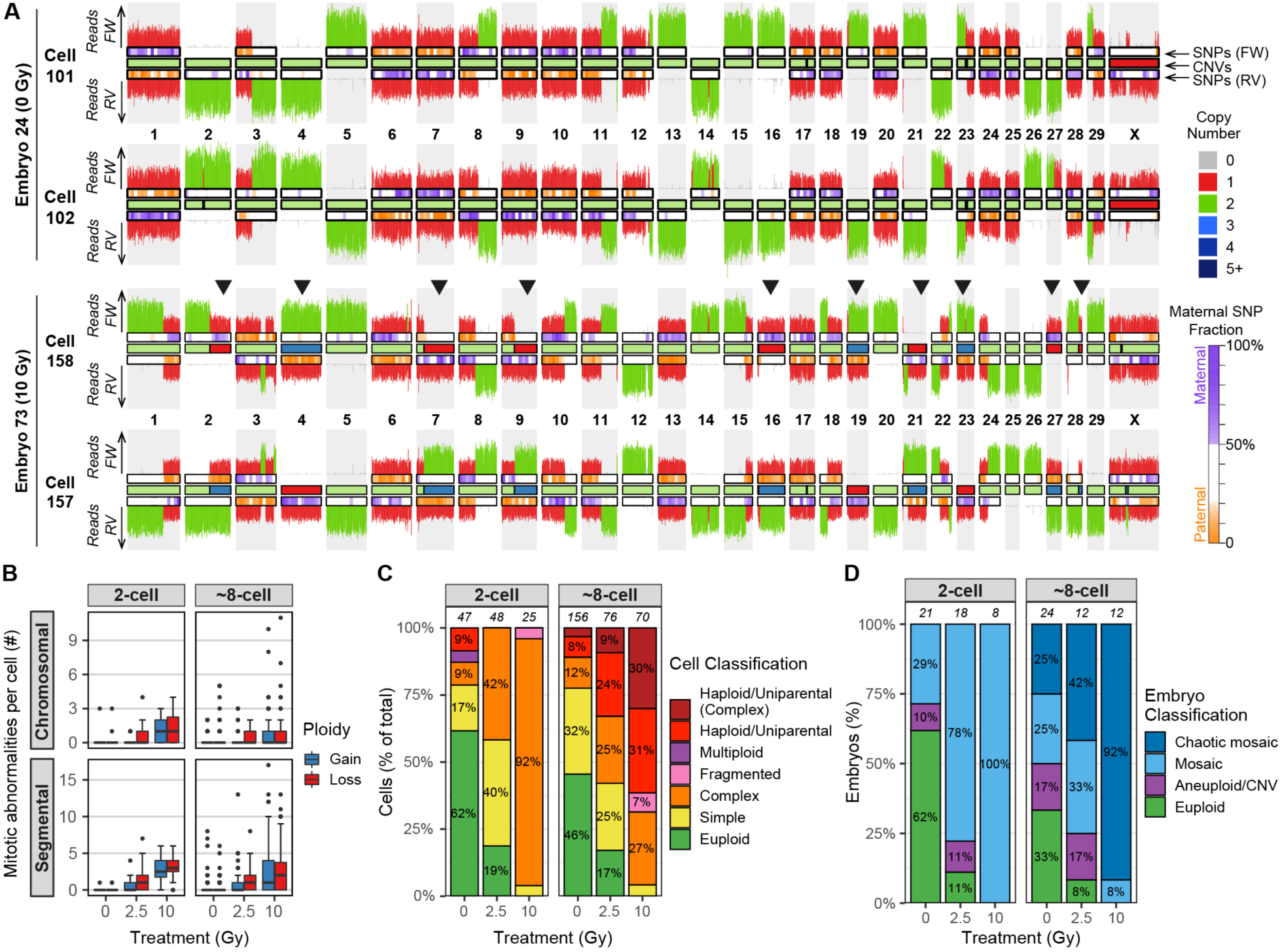
Sperm DNA damage causes mitotic errors and mosaicism in embryos. A) Representative examples showing chromosome ideograms with Strand-seq copy number profiles. Top: two-cell diploid control embryo; bottom: two-cell mosaic embryo produced with 10Gy-treated sperm. Arrowheads highlight mitotic errors. FW = forward/Crick strand, RV = reverse/Watson strand. (B) A radiation dosage dependent increase in the number of whole chromosome and segmental gains and losses per cell in two- and eight-cell stage embryos. (C) Classification of all the cells for the different treatment groups. The proportion of cells with multiple genomic abnormalities increases with sperm radiation dose. Cells with three or more chromosomal or segmental abnormalities are classified as complex. Fragmented cells only contain a few chromosomal fragments. Numbers above the bars indicate the number of analyzed cells per group. (D) Classification of all the embryos for the different treatment groups. The majority of embryos derived from fertilization with damaged sperm are mosaic, containing more than a single genotype. Embryos containing cells with a mix of more than three different genotypes are considered chaotic mosaic. The number of analyzed embryos per group is indicated above the bars.

Read-depth based copy number analysis revealed few copy number aberrations in the two-cell embryos produced with untreated sperm (Fig. 1B). Consistent with previous reports (*21, 27*), ∼14% of embryos produced with untreated sperm contained one or more copy number change due to a meiotic error (Fig. S2) and ∼19% of control embryos showed defects due to mitotic errors (Fig. 1C, D). Strikingly, the majority of embryos produced with damaged sperm (∼88%) showed multiple whole chromosome and segmental gains and losses with the number of aberrations increasing in a dose-dependent manner (Fig. 1B). Chromosomes or chromosomal segments that were gained in one cell were frequently lost in its sister cell resulting in an average disomic copy number state in the embryo, a phenomenon we refer to as mirrored mosaicism (Fig. 1A, Fig. 2A).

**Fig. 2.**
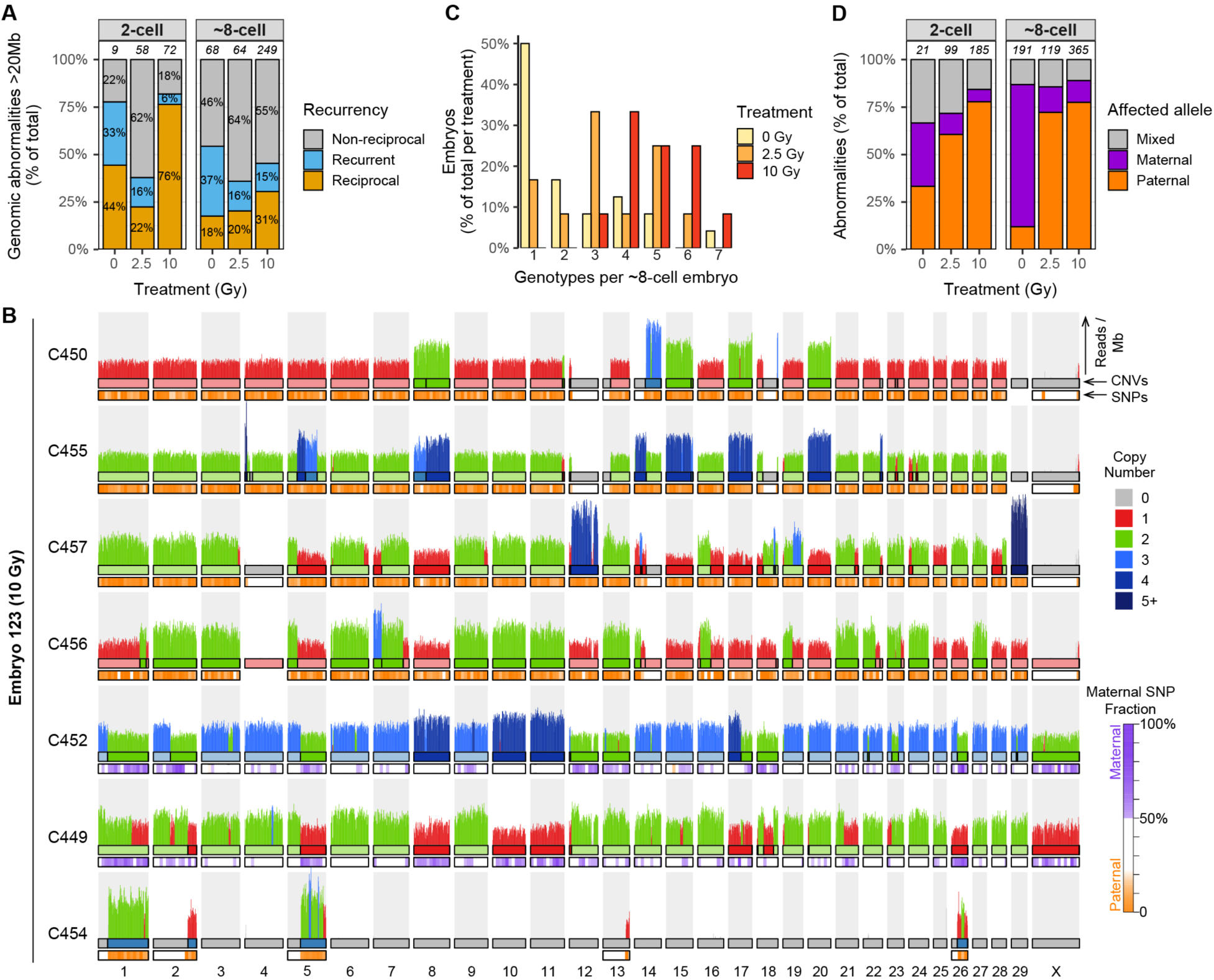
Chaotic mosaicism in eight-cell stage embryos produced with damaged sperm. (A) Many genomic abnormalities (>20 Mb in size) are mirrored between cells, showing reciprocal gains and losses between at least two cells. The relatively high number of non-reciprocal abnormalities, i.e. restricted to one cell per embryo, is largely due to variation in CNV calls between cells and in some cases due to missing or excluded cells. Numbers above the bars indicate the number of unique copy number variants per group. (B) Copy number profiles of seven sequenced cells from embryo E123 showing complex genomic abnormalities. Cells C450, C455, C456 and C457 are uniparental with only paternal chromosomes, whereas C449 and C452 are biparental. C454 is a fragmented cell containing chromosomal fragments that are complementary to copy number losses in C449 and C452. (C) Embryos produced with damaged sperm frequently show more than three different genetic lineages around the eight-cell stage of development, indicative of genomic instability through mitotic errors. (D) The majority of copy number changes (>10 Mb) in embryos derived from fertilization with damaged sperm are located on alleles inherited from the father. The copy number changes on the maternal alleles are largely caused by meiotic errors. Numbers above the bars indicate the number of analyzed copy number abnormalities per group.

To examine the genomic consequences at later stages we performed single-cell whole genome sequencing (*25*) of individual blastomeres at the ∼eight-cell stage of development (∼48 hpf). In total, 302 individual blastomeres of 48 embryos, of which 24 were derived from fertilization with damaged sperm, were successfully sequenced (Fig. S1B, C, Table S1). Embryos derived from fertilization with irradiated sperm contained fewer euploid cells and more cells with complex rearrangements, i.e. affecting at least three chromosomes (Fig. 1C). Copy number alterations were frequently observed and many were either shared or, as observed in the two-cell stage embryos, mirrored between blastomeres from the same embryo (Fig. 1B, Fig. 2A, B). For each condition, the average number of chromosomal aberrations per cell was similar between the two-cell stage and the eight-cell stage (Fig. 1B), which indicates no further fragmentation of chromosomes from the two-cell stage onwards. However, most eight-cell embryos derived from damaged sperm contain cells representing more than three different genotypes, indicating progressive genomic instability and segregation defects after the two-cell stage (Fig. 2B, C).

Bulk whole genome sequencing of the sperm DNA enabled haplotyping of the embryonic single cell sequencing data and revealed that copy number alterations were, as expected, strongly biased towards the paternally-derived chromosomes in both two and eight-cell stage embryos (Fig. 2D, Fig. S3), except for meiotic errors, which were biased towards maternal chromosomes (Fig. S2). These results indicate that post-meiotic sperm DNA damage results in fragmentation of the paternal genome followed by distribution of the DNA fragments over both daughter cells during the first embryonic cell division.

Chaotic mosaicism, where blastomeres of the same embryo have seemingly random chromosome complements, was common in embryos produced with irradiated sperm (Fig. 1D). The variety of genomic abnormalities ranging from aneuploidies, segmental changes, abnormal ploidy states, to cells containing minimal chromosomal content restricted to a few chromosomal fragments (Fig. 2B, Fig. S4), covers the broad spectrum of chromosomal aberrations that have been previously described in human, primate, and bovine embryos (*7, 21, 27, 28*). To further investigate the processes that contribute to chaotic mosaicism we performed Strand-seq on individual blastomeres of twelve ∼eight-cell stage embryos produced with damaged sperm. Strand inheritance patterns enabled a lineage reconstruction of chaotically mosaic embryos. From the strand-inheritance patterns of two embryos we could deduce that seven cells were formed by direct unequal cleavage of both blastomeres of a two-cell stage embryo that cleaved into three and four cells respectively (Fig. 3A, B, Fig. S5A). These observations indicate that sperm DNA damage can cause aberrant cleavage divisions at the two-cell stage embryo resulting in chaotic mosaicism at later stages. Notably, in a recent study in rhesus macaque embryos, chaotic aneuploidy was correlated with one particular sperm donor (*27*), which may indicate that sperm DNA damage is the underlying cause. Chaotic mosaicism is also common in human embryos produced with sperm from men with non-obstructive azoospermia, a condition that is also associated with high levels of sperm DNA damage (*29*). Complex abnormal mosaic embryos have reduced implantation and clinical pregnancy rates and reduced chances to develop to term (*30*). Chaotic mosaicism thus appears to be the responsible intermediate step for the well-established correlation between sperm DNA damage and reduced fertility (*31, 32*).

**Fig. 3.**
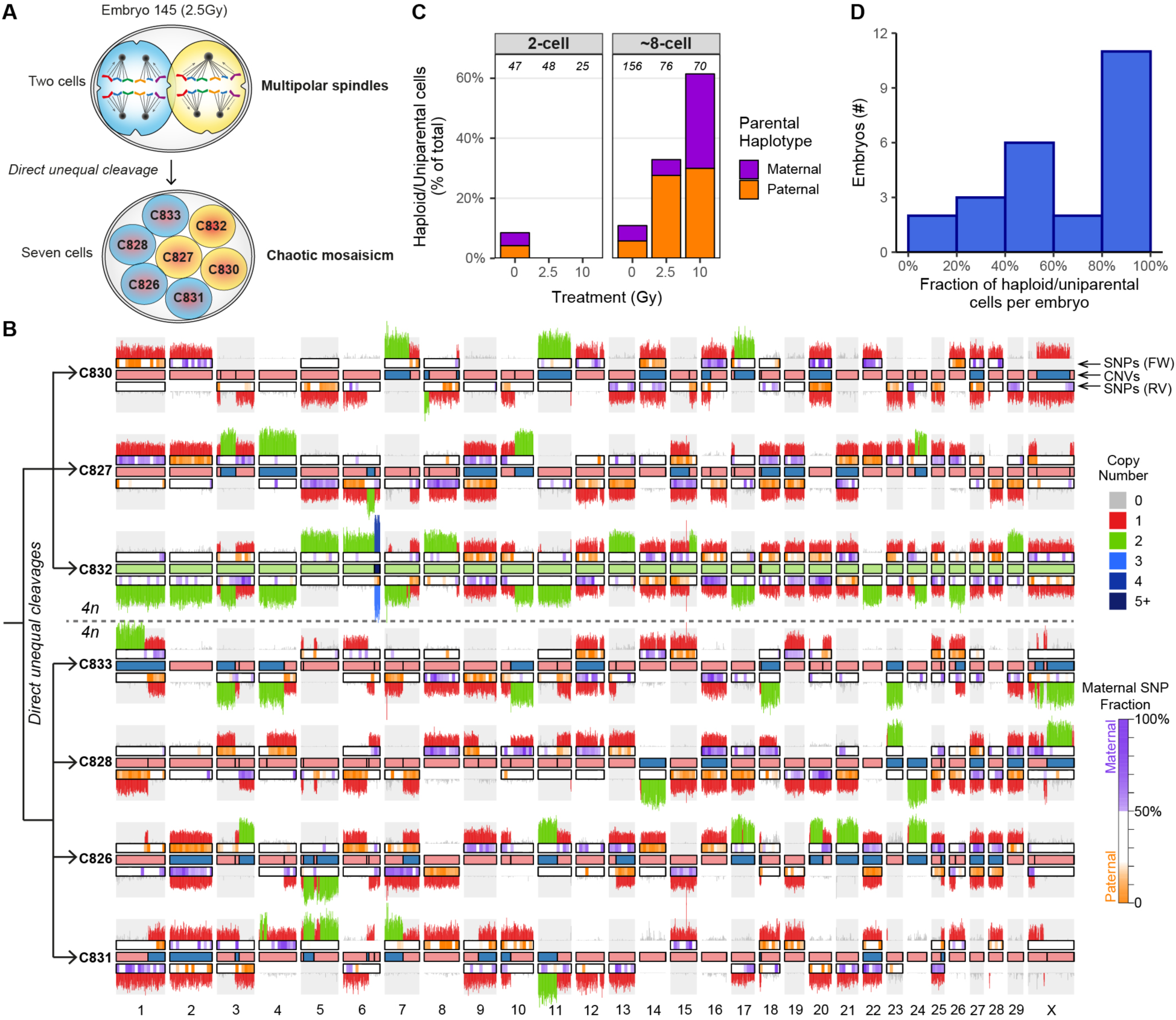
Multipolar cell divisions lead to chaotic mosaicism at the eight-cell stage. (A) Schematic reconstruction of the direct unequal cleavage divisions of two-cell stage blastomeres into respectively three and four daughter cells in embryo 170 (produced with 2.5 Gy-treated sperm). Sister cells are depicted with the same color. (B) Strand-seq karyogram of seven-cell embryo (E170) showing chaotic mosaicism after direct unequal cleavage divisions at the two-cell stage. The strand inheritance patterns detected by Strand-seq enable the identification of sister cells and the deduction of the preceding division and the distribution of the chromosomal fragments. This analysis reveals that both blastomeres at the two-cell stage performed a multipolar division; a tripolar division generated the sister cells C830, C827 and C832 and a tetrapolar division generated the sister cells C833, C828, C826 and C831. This led to the random distribution of the tetraploid set of chromosomes over the sister cells. As a consequence, the DNA fragments distributed over the sister cells sum up to a 4n copy number state. FW = forward/Crick strand, RV = reverse/Watson strand. (C) Embryos generated with damaged sperm contain more haploid and uniparental cells having a genomic content from either the father or the mother, indicating that sperm DNA damage causes heterogoneic cell divisions. Numbers above the bars indicate the number of analyzed cells per group. (D) In many embryos only half of the cells are haploid/uniparental, suggesting that in some cases haploid/uniparental cells may arise after the two-cell stage.

Strikingly, a large number of cells from eight-cell stage embryos produced with damaged sperm lacked X-chromosomes or contained nullisomies, indicating that these cells are (near) haploid or uniparental (Fig. 2B, Fig. S3C). To accurately quantify the number of haploid and uniparental cells, we screened for cells that lack heterozygous SNPs. Only a few cells are haploid and/or uniparental in two-cell stage embryos and in control eight-cell stage embryos (Fig. 1C, Fig. 3C). In contrast, the proportion of haploid/uniparental cells in eight-cell stage embryos produced with damaged sperm increased with radiation dose, amounting to two-thirds of the cells in the embryos produced with 10Gy-treated sperm (Fig. 1C, Fig. 3C).

A recent study described complete segregation of maternal and paternal genomes through so-called heterogoneic cell divisions, which were hypothesized to be the result of direct unequal cleavage of the zygote (*21*). To examine if this process can indeed lead to heterogoneic cell divisions, we sequenced all blastomeres from nine embryos containing three cells that were formed after a direct unequal division of the zygote. In three out of nine three-cell embryos (two control embryos, one from irradiated sperm), all sequenced blastomeres were haploid (Fig. S5B). These observations indicate that unequal cleavages of zygotes can indeed lead to heterogoneic cell divisions yielding uniparental lineages, but the number seems insufficient to explain the high incidence of haploid and uniparental cells observed at the eight-cell stage in embryos produced with damaged sperm. Because parental genomes still occupy distinct territories at the two-cell stage (*33-35*), uniparental and haploid cells may also be formed by unequal cleavages at this stage of development. The observation that frequently only half of the cells of eight-cell stage embryos were haploid/uniparental (Figure 3D) is indeed in support of heterogoneic cell divisions of two-cell stage blastomeres. Direct unequal cleavage divisions are the result of multipolar spindles. Strikingly, fertilized oocytes that carry the minor allele of *PLK4* are also vulnerable to multipolar spindle formation and mosaicism in embryos (*16, 17*). Thus, spindle aberrations appear to be an important source of genomic instability in embryo development and we here identify sperm DNA damage as one possible cause.

Gamma radiation results in fewer than 40 double-strand breaks per diploid human cell per Gy (*36*), causing at most 100 double strand breaks per cell when treated with 2.5 Gy. For haploid bovine sperm cells, the number of induced breaks per cell is presumably less. The comet assay is considered the most sensitive method to detect sperm DNA damage with an estimated lower bound of 100 double-strand breaks per cell (*37*). Our results from the 2.5 Gy embryos demonstrate that even upon the induction of sperm DNA damage close to or below this detection limit leads to embryonic genome instability. Since it is inherently impossible to know the degree of DNA damage of the individual sperm cell that was used to fertilize an oocyte, fertilizations with damaged sperm may be an underestimated phenomenon contributing to the widespread genomic instability in human embryos.

## Acknowledgments

We thank Wigard Kloosterman for helpful discussions and Roel Janssen for bioinformatics support. We would also like to thank the Hartwig Medical Foundation for whole genome sequencing.

## Author contributions

HvT and EK performed wet-lab experiments. DS, VG, and PL performed single cell sequencing. SM, HvT, DS, and EK performed data analysis. SM, HvT, DS, BR, PL, EC, and EK were involved in the conceptual design of the study. PL, EC, and EK acquired financial support for the study. SM, EC, and EK wrote the manuscript with input from all authors.

## Data and materials availability

All sequencing data have been deposited in the European Nucleotide Archive (https://www.ebi.ac.uk/ena) under accession number PRJEB32696. Custom code used in this study is available on GitHub (https://github.com/UMCUGenetics/Bovine_Embryo/).

## Supplementary Materials for

### Materials and Methods

#### Bovine IVF and blastomere collection

Fertilization and embryo culture were performed, according to previously described procedures (*38*). In short, bovine cumulus oocyte complexes were aspirated from 2-8mm antral follicles of ovaries that were obtained from the slaughterhouse. Subsequently, germinal vesicle stage oocytes with an intact multilayered cumulus were selected and matured in M199 supplemented with 26.2 mM NaHCO3, 0.05 IU/ml recombinant human FSH (Organon, Oss, The Netherlands), 10% Fetal Bovine Serum (Gibco BRL) and 1% (v/v) penicillin-streptomycin (Gibco BRL) at 39°C in a humidified atmosphere of 5% CO_2_ in air. In vitro fertilization was performed at 23 h after maturation with 0.5 × 10^6^ sperm cells per ml sperm. To obtain sperm with damaged DNA, sperm straws were subjected to ionizing radiation from a Gammacell 1000 (Atomic Energy of Canada Limited, Mississauga, Southern Ontario, Canada) prior to IVF. Ionizing radiation allows induction of DNA damage on non-cycling sperm cells while maintaining accurate control over the dosage. Untreated sperm from the same bull was used for the control group. All experiments were performed with sperm from the same donor bull to control for the potential natural variation in DNA damage between individuals. At 18–22 h after sperm addition, the cumulus cells and adhering sperm cells were removed and the denuded zygotes were further cultured in synthetic oviductal fluid (SOF) in a humidified incubator at 39°C with 5% CO_2_ and 7% O_2_. To obtain blastocysts, cleaved embryos were transferred to fresh SOF at day 5 and cultured until day 8. For Strand-seq experiments at the two-cell stage, bromodeoxyuridine (BrdU) was added to the fertilization medium and the embryo culture medium from the start of the embryo culture. Blastomeres were collected from 28h after fertilization (hpf) onwards. For Strand-seq experiments at the eight-cell stage, four-cell stage embryos (at 29-33 hpf) were transferred to medium containing BrdU and cultured until the eight-cell stage (at 48 hpf) when individual blastomeres were collected. For single-cell whole genome sequencing of eight-cell stage embryos, embryos were cultured in medium without BrdU and blastomeres were collected from 48hpf onwards. To collect individual blastomeres, embryos were placed in a droplet of 0.1% protease in PBS with 0.05% polyvinyl alcohol. After the zona pellucida was dissolved, the embryos were transferred to a droplet of Trypsin EDTA to dissociate the blastomeres. Blastomeres were transferred to single wells from a 96-well plate containing 5μl cryoprotectant consisting of 50% PBS, 42.5% ProFreeze (Lonza), and 7.5% DMSO. Full plates were stored at - 80°C until further processing.

#### Single-cell genome sequencing and primary data processing

Strand-seq and single-cell whole genome sequencing libraries were generated as previously described by respectively Falconer et al (*24*) and van den Bos et al. (*25*). Libraries were pooled (192 libraries per rapid run flow cell lane) and sequenced on the Illumina HiSeq 2500 sequencing platform. Raw sequencing reads were mapped to the *Bos taurus* UMD3.1 (bt8) reference genome using Bowtie2 (*39*) and BamUtil was used to filter duplicated reads. The median read count was 692,678 reads per cell after primary data processing (Table S1).

#### Single cell copy number variant calling and filtering

The BAM files for all single cell libraries were merged to generate a composite BAM file using Samtools merge. Bedtools intersect was used to calculate the coverage per 100kb genomic bins. A blacklist for CNV calling (included with the scripts) was generated by selecting the 3% bins with the highest and 2% of the bins with the lowest read counts on the autosomes and the bins with the top 5% and bottom 3% read counts on the X chromosome. The R-package AneuFinder (v1.8.0) was used to count the reads (with a minimal mapping quality of 10) in fixed-width bins of 1Mb and to call copy number variants using the “edivisive” method (36). The genomic sequence provided by the R-package BSgenome.Btaurus.UCSC.bosTau8 (v1.4.2) was used for GC-correction applied by Aneufinder. CNV calls with a limited change in read count compared to the median read count per bin per cell were excluded (decrease of <25% for presumed losses and increase of <25% for gains). Subsequently, CNV calls for each cell were merged based on a variable overlap threshold dependent on CNV size into one CNV call set per embryo (e.g. larger CNV require a higher percentage of overlap to merge than smaller CNVs). CNV calls occurring in more than 15% of the high-quality control libraries (with more than 200,000 reads), which likely correspond to common population variants or reference genome artifacts, were removed from the call sets. CNVs were considered to be reciprocal if there is at least one gain and one loss at a genomic location in the embryo. Sequenced libraries with more than 100,000 reads, 10 or less filtered chromosomal or segmental abnormalities, less than 80 genomic segments detected by Aneufinder and, if applicable, alternating Watson/Crick strand inheritance patterns (whose mother cell incorporated BrdU during replication and underwent mitosis) were used as high-quality libraries for further analyses (Table S1).

#### Bulk whole genome sequencing of bovine sperm DNA

Sperm DNA was extracted with the guanidine thiocyanate method (*40*). A Covaris sonicator was used to shear the isolated DNA to fragments of 400-500 basepairs. Libraries for whole genome sequencing were prepared using the TruSeq DNA Nano Library Prep Kit (Illumina) according to the manufacturer’s protocol. Paired-end 2×150 basepair read whole genome sequencing (2×150 base-pair reads) was performed on an Illumina Hiseq X sequencer to a mean genome coverage depth of 34x. Reads were aligned to the *Bos taurus* UMD3.1 reference genome using BWA-0.7.5a with settings BWA-MEM -t 12 -c 100 -M -R (*41*). Reads were realigned with GATK IndelRealigner (*42*) and duplicate reads were flagged with Sambamba markdup (*43*).

#### SNP genotyping of sperm and blastomere DNA

All non-reference single nucleotide variants (SNVs) were called from the composite BAM file (containing all the reads from the sequenced single-cell libraries) using bcftools mpileup and bcftools call (*44*). All heterozygous SNVs with more than 2 reference and 2 alternative allele counts and with a maximum coverage depth of 50 were selected to generate a list of 2,626,948 embryonic single nucleotide polymorphisms (SNPs). Subsequently, the paternal sperm WGS data was genotyped for the embryonic SNP positions using bcftools. To enable classification of SNPs in single embryonic cells as maternal (non-paternal), only the SNP positions that are homozygous in the father (with a coverage depth between 10 and 75 in the sperm WGS data) were selected. All the single cells were genotyped for these 912,144 homozygous sperm SNP positions using bcftools. A SNP was classified as maternally-inherited if the genotype is different from the homozygous genotype in the father.

#### Determination of the ploidy status of single blastomeres

Haploid and uniparental cells were identified based on several parameters. First all cells were genotyped for the 2,626,948 variable embryonic SNP positions in the composite BAM file (see above). Cells were considered to be uniparental if less than 15% or more than 50% of the called SNPs in the cell were different from the homozygous SNPs in the father (Fig. S3A).

Additionally, haploid cells were detected by a loss of heterozygous SNP positions.

Haploid/uniparental cells with more than 3000 covered SNPs were required to have less than one heterozygous REF/ALT SNP (excluding SNPs overlapping copy number gains) per 1000 called homozygous SNPs. Strand-seq libraries of haploid cells were recognized by the absence of bins with reads on both the Watson and Crick strands (haploid cells should only contain reads on one strand per bin after Strand-seq). Cells classified as haploid/uniparental were considered to be haploid (with a copy number state of one) if the majority (>80%) of called copy number losses are nullisomies.

#### Classifications of individual blastomeres and embryos

Cells were classified based on their ploidy status and the number of segmental and whole chromosome copy number changes. Cells containing three or more segmental or whole chromosome abnormalities were classified as complex. Cells with more than 10,000 reads and more than 25% of their reads on a single chromosome were considered to be fragmented. To determine the presence of different genotypes within each embryo, copy number changes (>20Mb) were compared between cells. Cells sharing more than 75% of their CNVs are considered to be of the same genotype. Embryos containing more than one or more than three different genotypes are classified as respectively mosaic and chaotic mosaic.

---

**Fig. S1.**
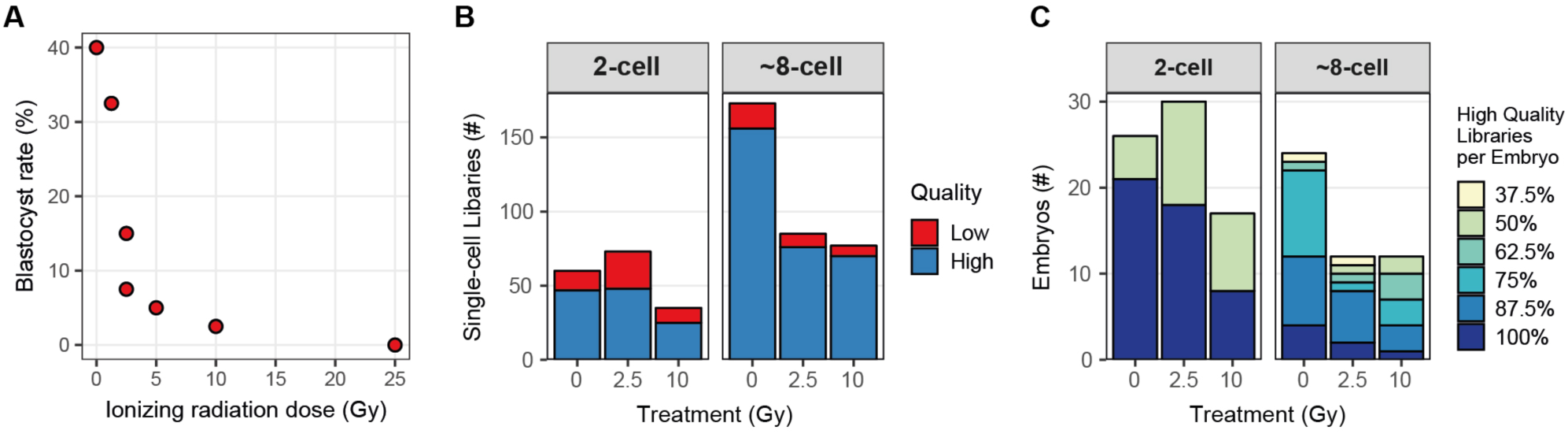
Characteristics of blastocysts and single cell libraries of analyzed bovine embryos. **(A)** Percentage of blastocysts that develop from fertilization with sperm treated with different doses of γ-radiation. Forty embryos per treatment group were produced with sperm radiated with 0, 1.25, 2.5, 5, 10, and 25 Gy. The number of blastocysts was counted at day 8 after fertilization. The results were obtained from two independent fertilization experiments. **(B)** Number of successfully sequenced single-cell libraries per developmental stage. High quality libraries have more than 100,000 non-duplicate reads with a mapping quality of more than 10 and 10 or less filtered chromosomal or segmental abnormalities. Strand-seq libraries additionally required the typical strand inheritance patterns. Sequencing results for low quality libraries with more than 10,000 reads are also included in the karyograms, because they can be informative for identifying sister cells, but they are excluded for further quantitative analyses. **(C)** Percentage of sequenced high-quality single cell libraries per embryo.

**Fig. S2.**
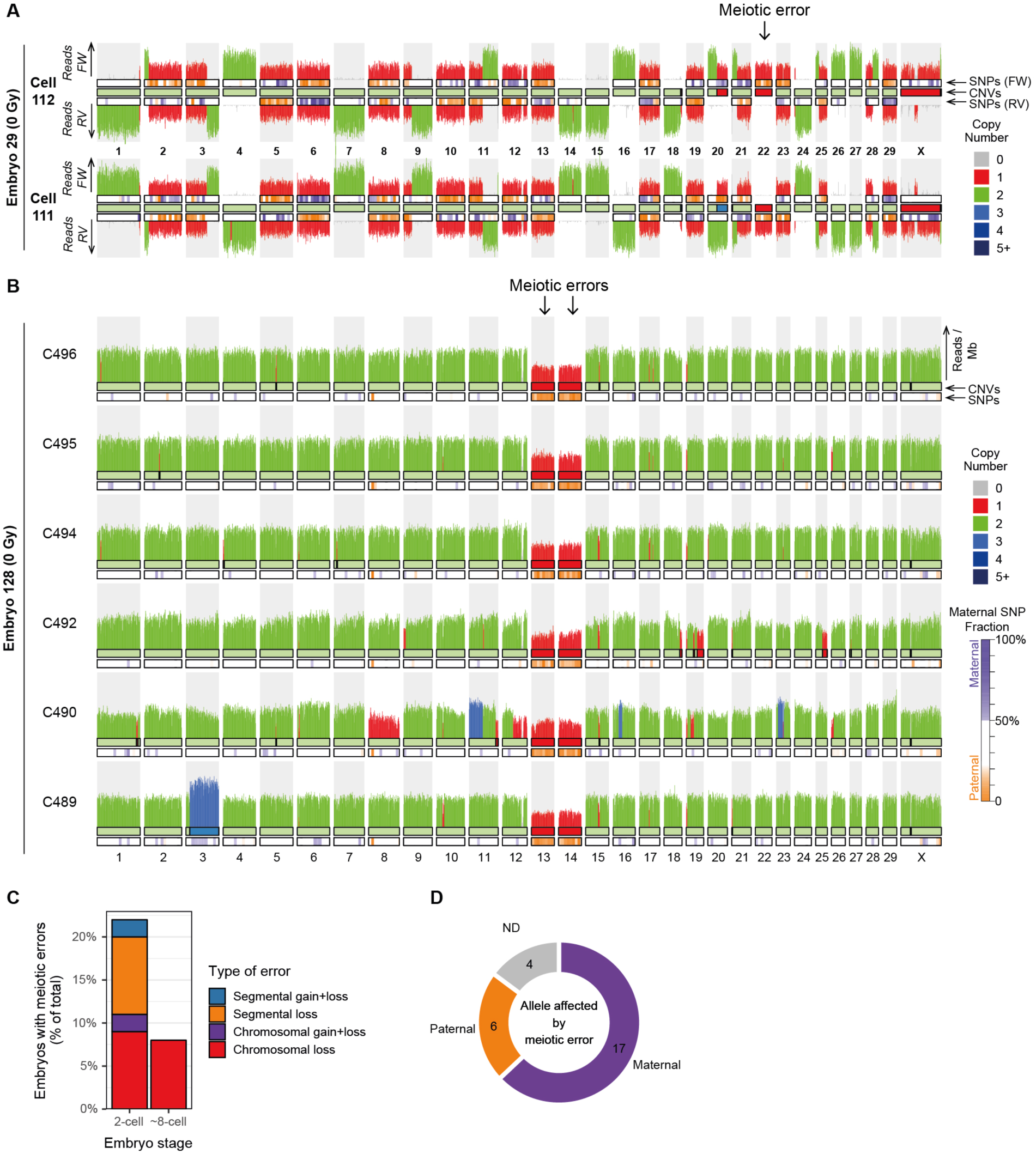
Incidence and parental origin of meiotic errors in early embryos. **(A)** Example of a two-cell embryo (E29) analyzed with Strand-seq containing a loss of chromosome 22 due to a meiotic error in the maternal germline. The remaining chromosome 22 is enriched for paternal SNPs. The embryo also contains a reciprocal mitotic copy number change on chromosome 20. FW = forward/Crick strand, RV = reverse/Watson strand. **(B)** Karyogram of six sequenced cells from one embryo (E133) showing meiotic losses of chromosomes 13 and 14. Only the paternally-inherited copies of chromosome 13 and 14 are present, indicating that the meiotic errors occurred on the maternal alleles. **(C)** Quantification of the number of cells containing different classes of meiotic abnormalities (>10Mb) per embryonic stage. **(D)** Number of meiotic errors (>10Mb) on maternally and paternally-inherited chromosomes.

**Fig. S3.**
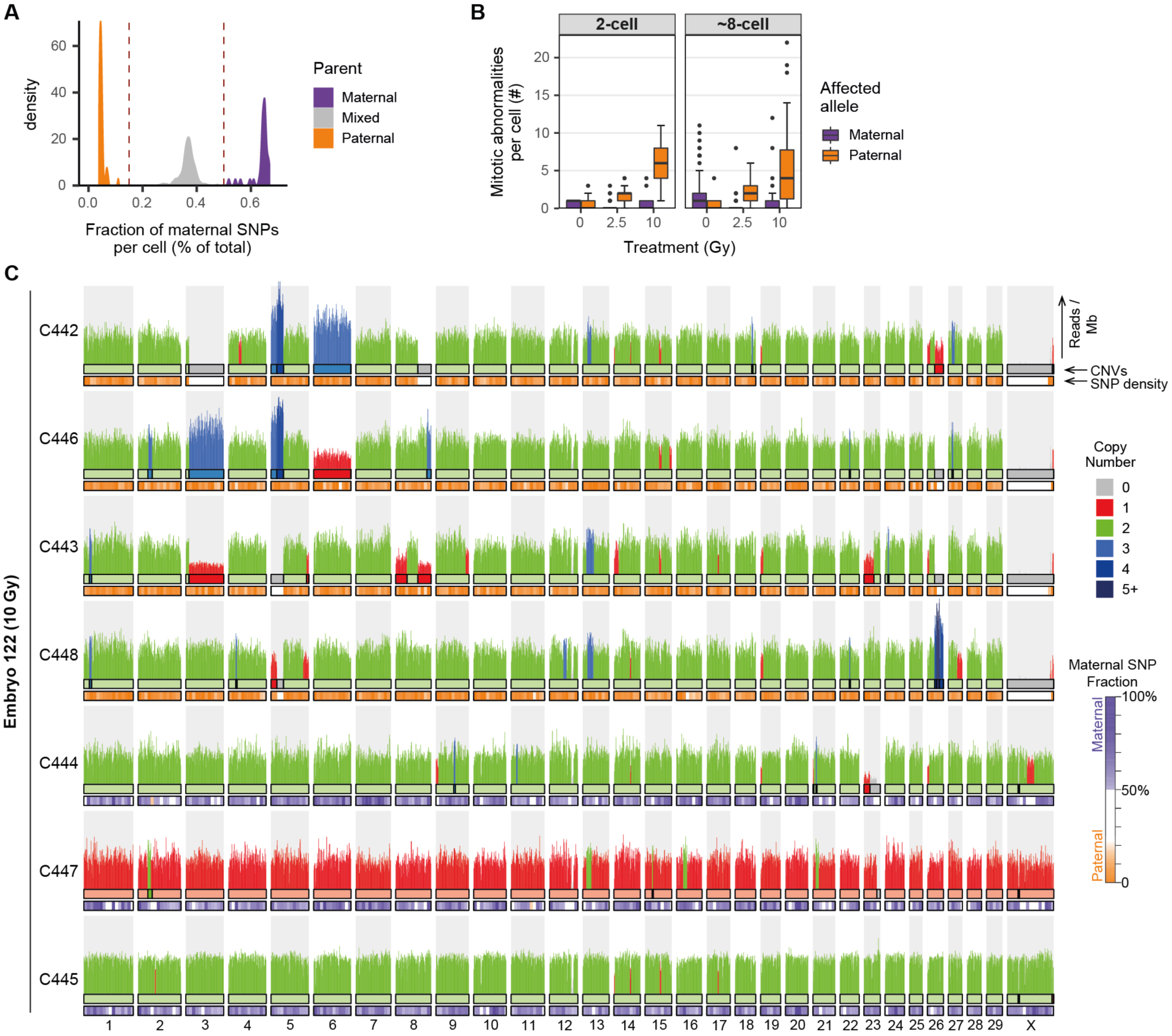
Bulk sperm DNA sequencing allows parental haplotyping of embryonic cells. **(A)** Proportion of SNPs in the single blastomeres corresponding to and diverging from homozygous SNPs in the genome of the father. Bulk whole genome sequencing enabled the detection of homozygous SNP positions in the genome of the bull whose sperm was used for all IVF experiments. SNPs in the blastomeres that are different from the SNPs in the father are considered to be maternally-inherited SNPs. SNPs overlapping between the father and the blastomeres can be paternally or maternally inherited (if the mother has the same SNP). **(B)** Sperm DNA damage leads to genomic abnormalities (>10Mb) on the paternally-inherited chromosomes. **(C)** Example karyogram of a seven-cell embryo (E122) produced with damaged sperm containing a segregation of uniparental maternal and paternal cells, suggesting a heterogoneic cell division of the zygote. Copy number changes are mostly present in the cells containing a paternal genome (C442, C443, C446 and C448) and cells with maternally-inherited chromosomes are relatively unaffected (although there is a loss of chr23 in cell C444).

**Fig. S4.**
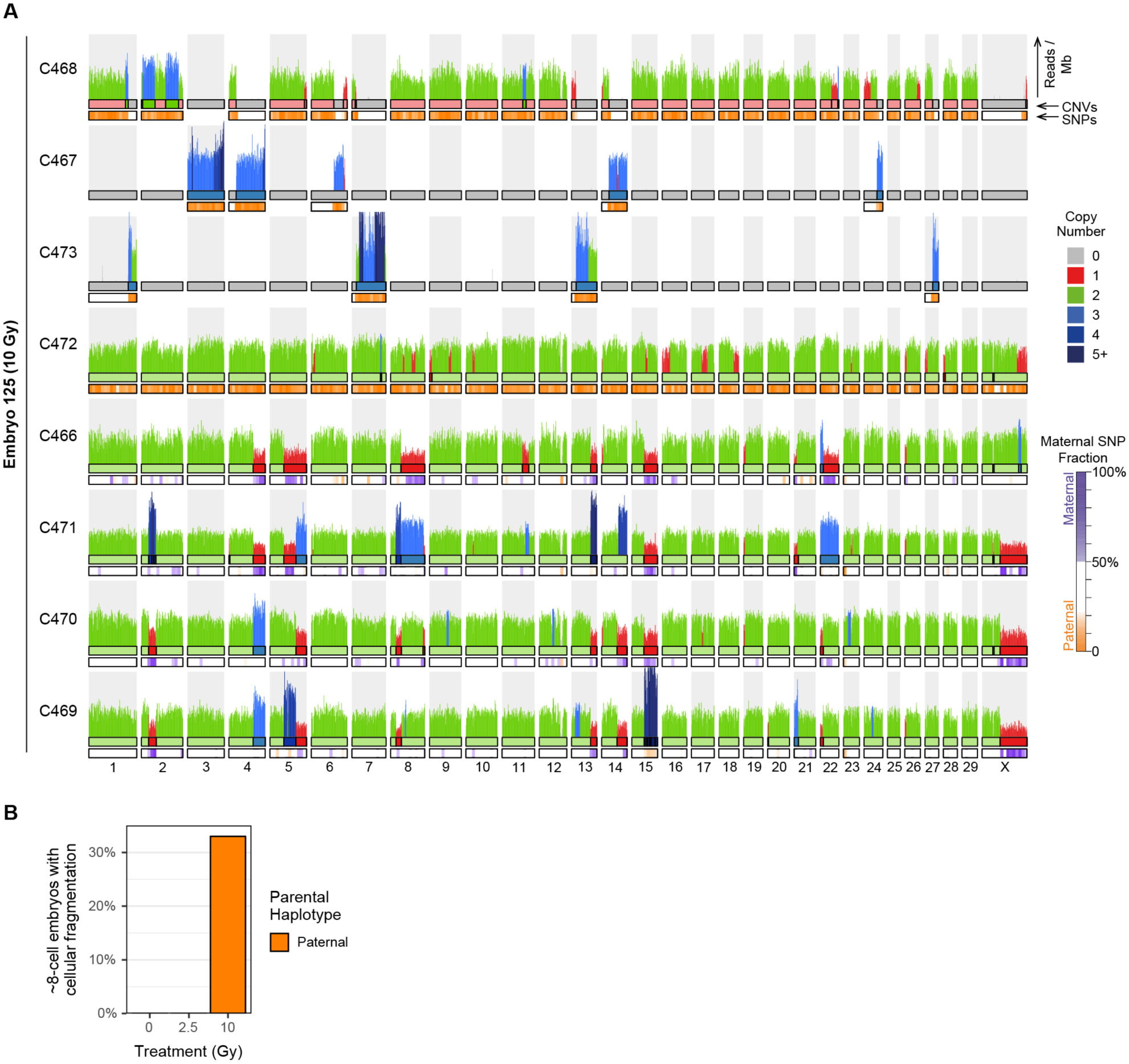
Cellular fragmentation is common in embryos derived from damaged sperm. **(A)** Example of an eight-cell stage embryo (E125) produced with damaged sperm (10Gy) showing cellular fragmentation. The mother cell of C468 is fragmented into three cells of which two cells (C467 and C473) only contain a few (respectively 5 and 4) chromosomal fragments. The top four cells only contain paternally inherited chromosomes. Cells C466, C469, C470 and C471 are diploid, but show many segmental gains and losses due to mitotic errors. **(B)** Quantification of the eight-cell stage embryos containing fragmented cells. Around a third of the eight-cell stage embryos produced with 10Gy-treated sperm contain fragmented cells with only paternally inherited chromosomes.

**Fig. S5.**
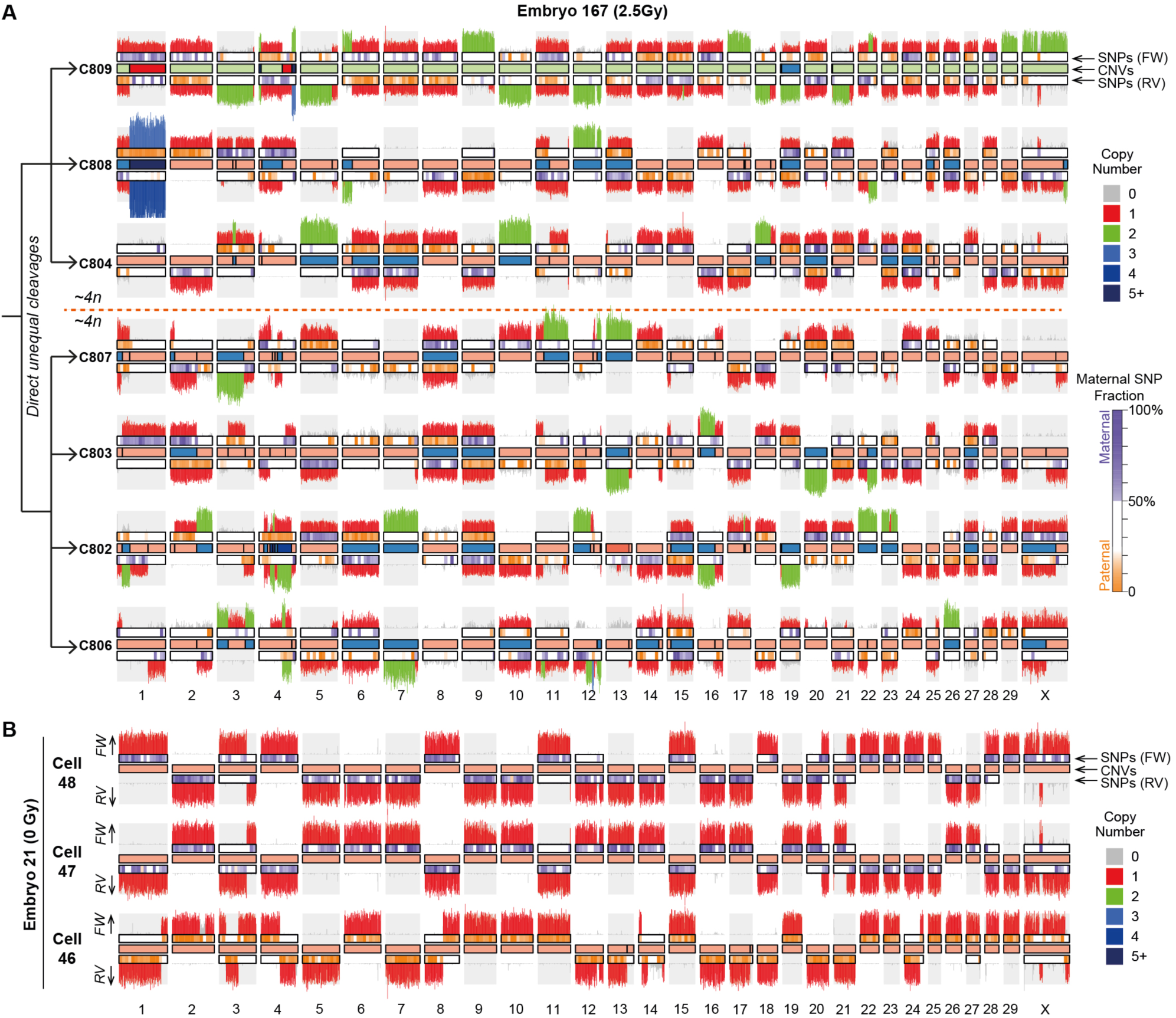
Direct unequal cell divisions at zygote and two-cell stage cause complex genomic rearrangements. **(A)** Karyogram of a seven-cell embryo (E167) analyzed by Strand-seq showing the results of direct unequal cell divisions at the two-cell stage. The strand inheritance patterns indicate that cells C809, C808 and C804 are sister cells as are cells C807, C803, C802 and C806. The DNA fragments distributed over the sister cells sum up to a 4n copy number state. **(B)** Example of three haploid cells originating from a heterogoneic cell division at the zygote stage, which lead to segregation of the paternal and maternal genomes. Sister cells C47 and C48 contain a haploid maternal genome. The sister cell of C46 may have been lost during collection of the single blastomeres. FW = forward/Crick strand, RV = reverse/Watson strand.

**Table S1.**

Summary for each analyzed blastomere. Negative/positive control cells are not listed in this table, resulting in fewer rows than cell numbers.

